# The spatial landscape of infectious mononucleosis tonsils

**DOI:** 10.1101/2025.10.09.681345

**Authors:** Ciara I. Leahy, Matthew R. Pugh, Aoife Hennessy, Nadezhda Nikulina, Stefan Dojcinov, Gerald Niedobitek, Graham S. Taylor, Paul G. Murray, Éanna Fennell

**Author notes:** Correspondence: Éanna Fennell & Paul G. Murray (;).

## Abstract

Primary Epstein-Barr virus (EBV) infection remains incompletely understood, particularly in relation to the nature of *in vivo* latency patterns and their influence on the immune microenvironment. Here, we present the first multiomic single-cell spatial atlas of palatine tonsils from individuals with infectious mononucleosis (IM). We identify rare EBV infection of epithelial cells, challenging the prevailing assumption that primary infection is restricted to B- and T-cells. Moreover, our detailed analysis suggests a spatiotemporal transition of EBV latency programs, with distinct viral gene expression patterns corresponding to specific tissue microenvironments-a dynamic process previously inferred but not directly mapped in human tissues. We found that cells expressing the EBV-encoded protein LMP1 form an immunosuppressive niche that is rich in IDO1+PDL1+ macrophages and relatively depleted of T-cells. Furthermore, the T-cells that do infiltrate LMP1+ regions exhibit signatures consistent with an activated and cytotoxic phenotype, providing new insights into how LMP1 shapes immune evasion. Taken together, our data redefine the spatial and functional organisation of primary EBV infection, integrating latency plasticity, epithelial tropism, and microenvironmental immunosuppression into a unified framework, as well as providing a data-rich framework for understanding viral persistence and immune evasion.

## Introduction

The Epstein-Barr virus (EBV) infects 95% of the global adult population^1^. While most infected individuals carry EBV asymptomatically for life, the virus contributes to the pathogenesis of multiple cancers, including B-cell lymphomas and epithelial cancers^2^, as well as autoimmune diseases, such as multiple sclerosis (MS)^3^. Primary EBV infection usually occurs in infancy and is asymptomatic ^4^. However, if infection is delayed until early adulthood it can result in a clinical syndrome known as infectious mononucleosis (IM), which is characterised by fever, lymphadenopathy and fatigue^5^. Although IM is almost always a self-limiting benign disease^6^, a prior history of IM is associated with a 4-fold increased risk of developing EBV-associated classical Hodgkin lymphoma (cHL), a 7.1-fold increased risk of nasopharyngeal carcinoma (NPC) and a 2.3-fold higher risk of developing MS, in the context of certain MHC haplotypes and increased EBNA1 titres^3,7,8^. Thus, a better understanding of the pathogenesis of IM could provide insights into the nature of primary EBV infection and alterations of EBV innate and adaptive immunity that may contribute to the development of EBV-driven malignancy and autoimmune disease.

Despite its relevance to the development of viral driven illnesses, the spatial immunobiology of primary EBV infection is only poorly understood. Models to explain colonisation of the B cell pool during the establishment of persistence are largely based on studies of EBV infection *in vitro* and RT-PCR analysis of the blood and tissue of healthy EBV carriers and IM sufferers^9,10^. The prevailing consensus is that EBV initially infects naïve B-cells, which then enter a so-called latency III program characterised by the expression of six viral proteins located in the nucleus of infected cells (known as Epstein–Barr nuclear antigens; EBNAs-1, -2, -3A, -3B, -3C, and - Leader Protein; LP), and three viral proteins present in the plasma membrane (known as latent membrane proteins; LMP1, LMP2A, and LMP2B)^11,12^. This is thought to be followed by a germinal centre (GC)-like phase in which only EBNA1 and the LMPs are expressed, referred to as latency II or the ‘default program’. During this phase, LMP1 and LMP2A, which are functional mimics of CD40 and the B-cell receptor (BCR), respectively, enable the survival of EBV-infected cells and their exit from the GC reaction, eventually becoming memory B-cells^13^. EBV-infected memory B-cells almost entirely shut down viral gene expression (known as latency 0) but can activate EBNA1 expression to facilitate the replication and segregation of viral episomes during memory B-cell proliferation (latency I)^14,15^. These EBV-infected memory B-cells can also differentiate into plasma cells, a process that is accompanied by the induction of the viral lytic cycle^16^.

Although we have a detailed understanding of the nature and regulation of different EBV latency states in cells infected *in vitro*, significant gaps remain in our understanding of primary infection *in vivo*. For instance, it is unclear how EBV regulates latency within the context of a dynamic and reactive immune microenvironment. There is also limited knowledge about the spatial organisation of EBV-infected cells within these tissues and how the physical proximity of infected cells to immune effector cells impacts disease. Moreover, the mechanisms by which EBV-infected cells migrate to and persist within specific tissue niches, including the epithelial cells of palatine tonsils, proposed to be a site of viral transfer into and out of the oropharynx, are poorly defined. Indeed, a role for epithelial cells in primary infection, remains controversial. Thus, while rare EBV-positive epithelial cells have been detected in the tonsils of asymptomatic EBV-positive individuals and in the tongue tissues of immunosuppressed individuals^17^, EBV infection of epithelial cells in IM has not previously been demonstrated. Addressing these unknowns is essential for a comprehensive understanding of EBV pathogenesis and its long-term sequelae, including the development of cancers and autoimmune diseases.

Integrating spatial transcriptomics and multiplex immunofluorescence analysis of >1.2M cells, we have created the first single-cell spatial atlas of palatine tonsils from people with IM, revealing three key advances: (1) rare EBV infection of epithelial cells-a cell type excluded from canonical models of primary infection, challenging historical paradigms; (2) spatiotemporal mapping of EBV latency programs, showing microenvironment-associated viral gene expression gradients that correlate with immune cell exclusion; and (3) discovery of LMP1+ B-cell niches that recruit immunosuppressive IDO1+PDL1+ macrophages, deplete T cells and induce T-cell activation. By linking these findings to tonsillar immune architectures, we resolve longstanding debates about EBV’s cellular tropism and immune evasion strategies while providing a resource (latency maps, LMP1 interactome) for studying EBV-associated malignancies.

## Results

### A multiomic single-cell spatial atlas of IM tonsil reveals EBV infection of epithelial cells

Archival surplus to diagnostic requirement was available from formalin-fixed paraffin-embedded (FFPE) tissues of palatine tonsils from eight patients excised during the active phase of IM, typically 4-6 weeks post primary EBV infection^18^. A clinical and morphological diagnosis of infectious mononucleosis was established and reviewed (by consultant pathologists SD & GN). EBV infection was confirmed by EBER *in situ* hybridisation (ISH) staining, showing variable positive staining in IM tissues (Extended Data Fig. 1a-b). EBER ISH, along with Haematoxylin and Eosin (H&E) stains, were used to identify regions of active infection, exclude necrotic areas and select regions to generate tissue microarrays (TMA; Extended Data Fig. 1c). TMAs were subject to an OPAL-based mIF panel, allowing marker detection across six channels (OPAL mIF; n = 8), developed to detect the spatial co-localisation of the EBV latent proteins EBNA1, EBNA2, LMP1, LMP2A and lytic cycle induction protein, BZLF1 (Fig. 1b and Supplementary Table 1), a 1000-plex CosMx spatial transcriptomics (ST; n = 4) panel, and multiplex immunofluorescence with the PhenoCycler-Fusion (PF mIF; n = 8) (Fig. 1a and Extended Data Fig. 1d). A 42-plex PF mIF panel targeting cellular proteins and the viral proteins EBNA1, EBNA2, and LMP1 was utilised to identify EBV-infected cell types and a separate 50-plex PF mIF panel was employed to evaluate the cellular response to EBV infection (Fig. 1c & Supplementary Table 2 & 3).

**Figure 1.**
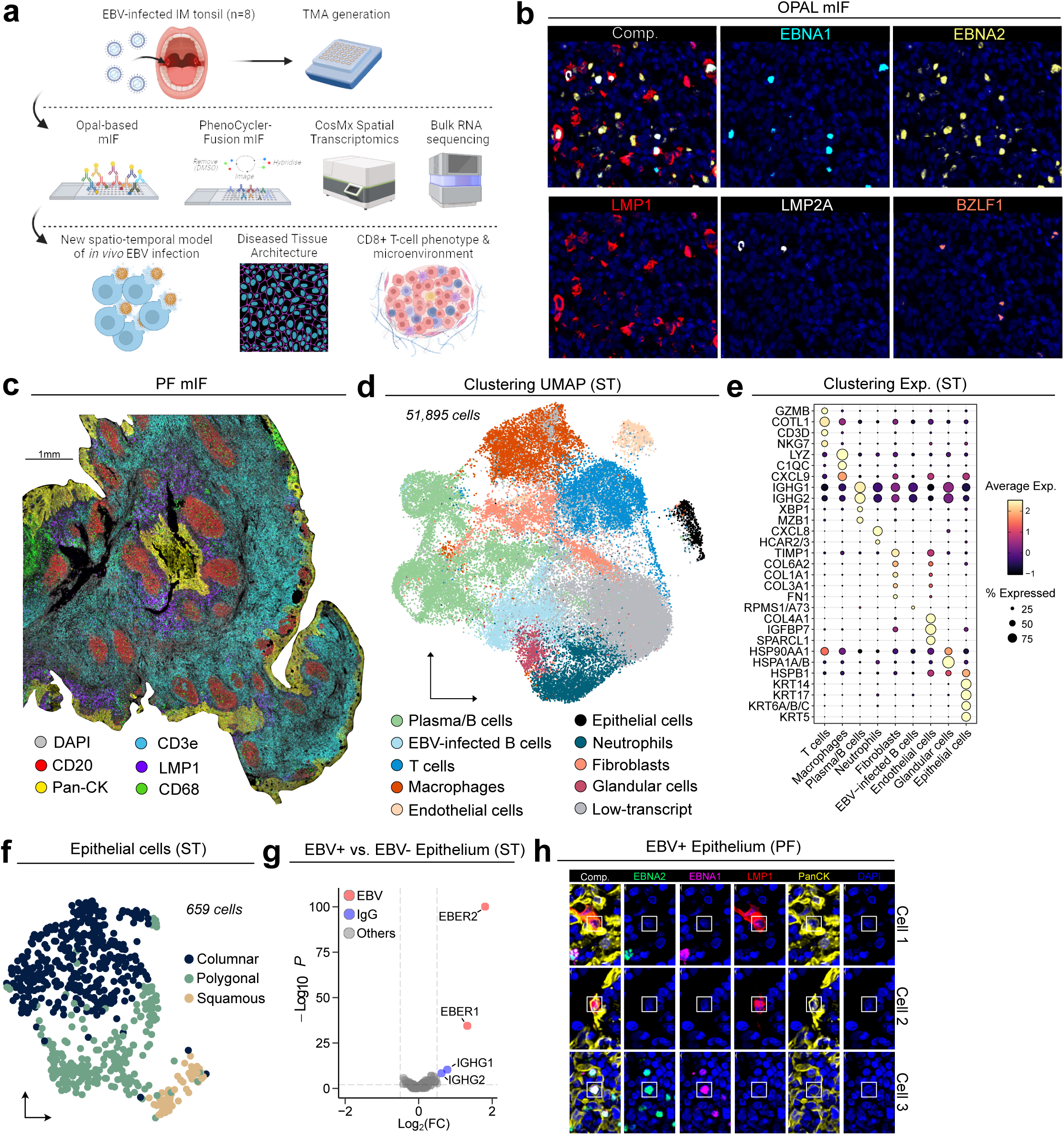
| A multiomic single-cell spatial atlas of infectious mononucleosis. **a.** Schematic overview of the multiomic spatial single-cell workflow. **b.** OPAL mIF staining of developed and validated EBV-panel showing co-expression of EBV viral latent antigens EBNA1, EBNA2, LMP1 & LMP2A as well as lytic protein BZLF1. **c.** PhenoCycler-FUSION mIF staining of full-face IM tonsil tissue showing human CD20 (red), pan-cytokeratin (yellow), CD3e (blue), CD68 (green) and the EBV viral protein LMP1 (purple). **d.** Uniform manifold approximation and projection (UMAP) of all 51,895 cells captured by CosMx spatial transcriptomics (ST), coloured and labelled by the 9 populations identified from Louvain single-cell clustering. **e.** Dot plot showing the average gene expression of the top differentially expressed genes per cluster, canonical lineage markers as well as EBV genes for the CosMx ST annotated populations. **f.** UMAP of re-clustered epithelial cells from CosMx ST showing columnar (basal), polygonal (intermediate) and squamous (superficial) phenotypes. **g.** Differential gene expression analysis of EBV+ vs. EBV-epithelial cells highlighting EBV EBERs and IgG expression **h.** Representative images of EBV protein expression in the PF mIF data showing co-expression with pan-cytokeratin.

Image analysis including TMA de-arraying, normalisation, single cell segmentation and feature extraction was completed using CellMAPS and identified ∼1M cells (see Methods)^19^. PhenoGraph clustering assigned 15 broad cell types annotated by cellular expression profiles, and each cell was further assigned a binary positive/negative score for functional marker expression (Extended Data Fig. 1d-e; Supplementary Table 4). Complementary to this, we profiled 51,895 cells by CosMx with a 1000-plex panel spiked with 24 EBV RNA targeting probes (Supplementary Table 5)^20^. Louvain clustering of the CosMx single-cell data identified 9 broad cell types, matching those identified by PF mIF (Fig. 1d) which were annotated according to canonical cellular expression profiles (Fig. 1e).

Within the ST assay, although the expression of latent and lytic EBV genes was mostly constrained to B cells, including plasma cells, it was also seen in some T cells and in a small subset of epithelial cells (Extended Data Fig. 1f-g). Although it is widely accepted that EBV preferentially infects B cells and some T cells in IM^21^, whether EBV also infects epithelial cells during primary infection remains unresolved. Within the epithelial cell cluster, we identified three differentiation states: columnar-characterised by CD44 and KRT14 expression, polygonal-characterised by KRT13 and KRT14 expression, and squamous-characterised by KRT13 and KRT4 expression, representing basal, intermediate and superficial layers, respectively^22^ (Fig. 1f & Extended Data Fig. 1h). EBV latent and lytic antigens were present in all three states, but preferential infection of the squamous (superficial) layer was observed (Extended Data Fig. 1i-j). Differential expression analysis revealed increased expression of IGHG1 & IGHG2 (encoding IgG1 and IgG2 respectively) in EBV-infected epithelial cells compared to EBV-negative epithelial cells (Fig. 1g). Notably, no other B cell markers were expressed by these cells. We stained the tissues with the RP215 antibody, which detects non-B cell-derived sialylated IgG, which is associated with epithelial-to-mesenchymal transition (EMT)^23^, and observed strong staining of crypt epithelium in all cases (Extended Data Fig. 1k). Consistent with this, EBV infected epithelial cells expressed many EMT markers including GSN, LCN2 and SQSTM1 and downregulated PFN1 (Supplementary Table 6)^24–27^. A separate mIF assay for lineage markers and EBNA1, EBNA2 and LMP1 revealed expression of these EBV genes in pan-cytokeratin-positive cells and CD3-positive cells (Fig. 1h & Extended Data Fig. 1l-m). Together, these data demonstrate that in addition to infecting B and T cells, EBV infects epithelial cells during primary infection.

### Spatio-temporal evolution of B cell EBV-infection *in vivo*

We next analysed EBV protein expression using a stringently validated OPAL mIF panel (Extended Data Fig. 2a). Overall, across eight different tonsil samples, expression of either EBNA2 alone or of LMP1 alone were the most abundant patterns of expression, respectively (Fig. 2a & Extended Data Fig. 2b). Co-expression of LMP2A and the lytic switch protein BZLF1 was not observed (Extended Data Fig. 2b), re-enforcing a role for LMP2A in supressing lytic cycle induction in B cells *in vitro* ^28,29^. Moreover, a significant inverse correlation between the abundance of cells expressing EBNA2 or LMP1 was observed across the tonsil samples from different individuals, suggesting a dominance of these antigens at different stages of infection (Extended Data Fig. 2c). Cells co-expressing LMP1 and LMP2A were rare (∼1%) and were located outside germinal centres (Fig. 2b and Extended Data Fig. 2b). EBNA2-expressing cells were most often present among or immediately adjacent to epithelial cells, a subset of which also expressed BZLF1, potentially indicating early infection (Fig. 2b & Extended Data Fig. 2d). Using Moran’s I as an indicator of spatially clustering, cells expressing LMP1+ were most likely to co-locate, consistent with previous observations *in vitro* (Extended Data Fig. 2e).

**Figure 2.**
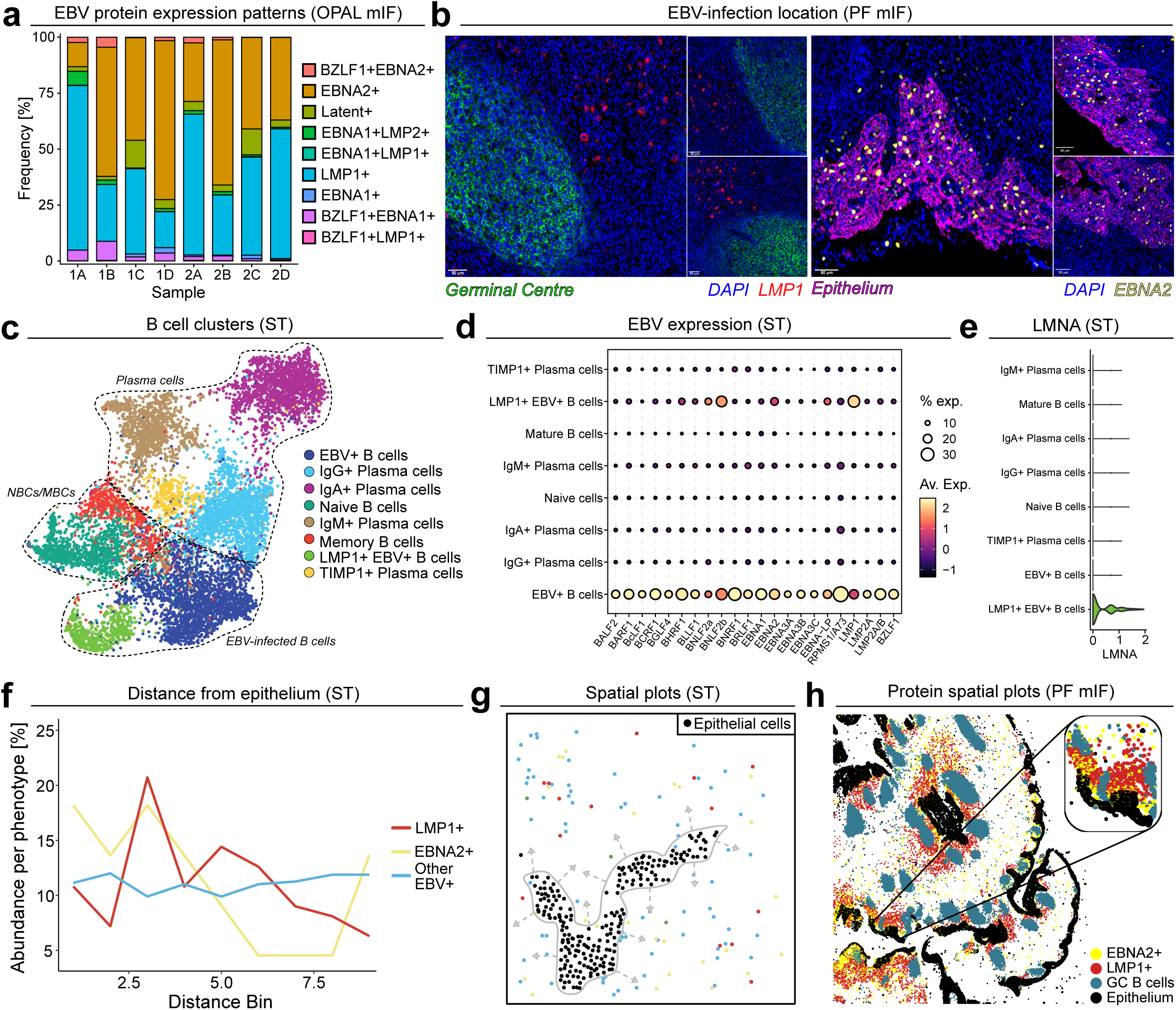
| Spatio-temporal evolution of EBV-infection in tonsillar tissue. **a.** Relative frequencies of different EBV protein co-expression patterns (OPAL mIF). Some co-expression patterns have been amalgamated for easier viewing; E1+ = EBNA1+, E2+ = EBNA2+, L1+ = LMP1+, L2+ LMP2+, B+ = BZLF1+, Latent+ = EBNA1+, EBNA2+, LMP1+, LMP2+ (see Supp. Table S11). EBNA2+ and LMP1+ constitute the main patterns. **b.** Localisation of LMP1 and EBNA2 protein expression to outside the germinal centres (CD21; green) and along epithelial beds (Pan-cytokertain; purple) respectively. **c.** UMAP of the re-clustered B and plasma cell lineage from ST data showing EBV+ clusters, naïve and memory B cells as well as IgM+, IgA+ and IgG+ plasma cells. **d.** EBV gene expression patterns within B and plasma cell clusters from ST data. **e.** Expression of human Lamin A (LMNA) in B and plasma cell clusters from ST data. **f.** LMP1+ (red), EBNA2+ (yellow) and other EBV+ (blue) B cell frequences as a function of distance from epithelial cell beds. LMP1+ and EBNA2+ cells were denoted by positive expression (>0) for LMP1 and EBNA2 respectively. **g.** Spatial plots of LMP1+ (red), EBNA2+ (yellow) other EBV+ cells (blue) and epithelial cells (black) showing the distances between epithelial beds and different EBV gene patterns from ST data. **h.** Spatial plots (PF mIF) confirming EBNA2+ and LMP1+ cell localisation at the protein level.

To gain more insight into the spatio-temporal dynamics of B cell EBV-infection, we re-clustered B cell phenotypes from CosMx ST, identifying naive and mature B cells as well as IgM+, IgG+ and IgA+ plasma cells (Fig. 2c & Extended Data Fig. 2f). Although EBV+ B cells were present in all differentiation states (defined by positivity for EBER1 & EBER2; Extended Data Fig. 2g), we identified two EBV+ clusters, seemingly independent of the standard B-cell differentiation pathway and exhibiting reduced expression of immunoglobulins and B cell markers (Extended Data Fig. 2f). The EBV+ clusters displayed different EBV expression profiles (Fig. 2d), with EBV cluster 1 expressing a range of lytic and latent EBV genes whereas EBV cluster 2 had low levels of lytic genes and high expression of LMP1 (Extended Data Fig. 2h). The LMP1+ EBV+ B cell cluster also expressed lamin A (LMNA), which is known to be regulated by LMP1 (Fig. 2e)^30^. To validate the observation that LMP1 and LMP2A were only rarely co-localised in the OPAL mIF dataset, co-expression of LMP1 and LMP2A/B was investigated in the ST assay (Extended Data Fig. 2I). This revealed that cells which were LMP1+ rarely expressed LMP2A/B, and vice versa. Furthermore, our spatial analysis also allowed us to plot EBV gene expression as a function of distance from epithelium (Fig. 2f). This revealed that EBNA2+ cells were most frequent around epithelial cells, while LMP1 positive cells were more frequently further from the epithelium (Fig. 2f-g). This spatial distribution of EBV+ cells was validated at the protein level, with EBNA2+ cells located immediately adjacent to the epithelium and LMP1 positive cells locating adjacent to these EBNA2+ cells (Fig. 2h & Extended Data Fig. 2j). Taken together, these data suggest a spatial organisation of EBV expression profiles within IM tonsillar tissue.

### LMP1-induced chemokines remodel the immune architecture of IM

Next, we investigated the cellular response to EBV infection, focusing on the phenotype of infected cells and the nature of their surrounding microenvironment. Analysis of the PF mIF dataset revealed that LMP1+ B cells downregulated the B cell marker CD20 compared to GC B cells, while CD44, PDL1, IDO1 and CD30 were upregulated compared to all other B cell subsets, in line with known LMP1 targets^31–33^ (Fig. 3a). Differential gene expression analysis of LMP1+ EBV+ B cells compared to EBV+ B cells by CosMx ST revealed the upregulation of other known LMP1 targets such as NFKBIA, CD40 and vimentin (Fig. 3b). Pathway analysis of genes upregulated in LMP1+EBV+ B cells revealed the enrichment of genes involved in cytokine responses, as well as inflammatory and interferon gamma associated response pathways (Fig. 3c). Accordingly, cytokines/chemokines and inflammatory genes upregulated in LMP1+EBV+ B cells compared to EBV+ B cells included macrophage attractants CXCL9, CXCL10 (encodes IP-10), macrophage/T cell activation cytokines, CCL18, CCL19 and CCL22 and MHC Class II genes, HLA-DRB and HLA-DPA1 (Fig. 3b).

**Figure 3.**
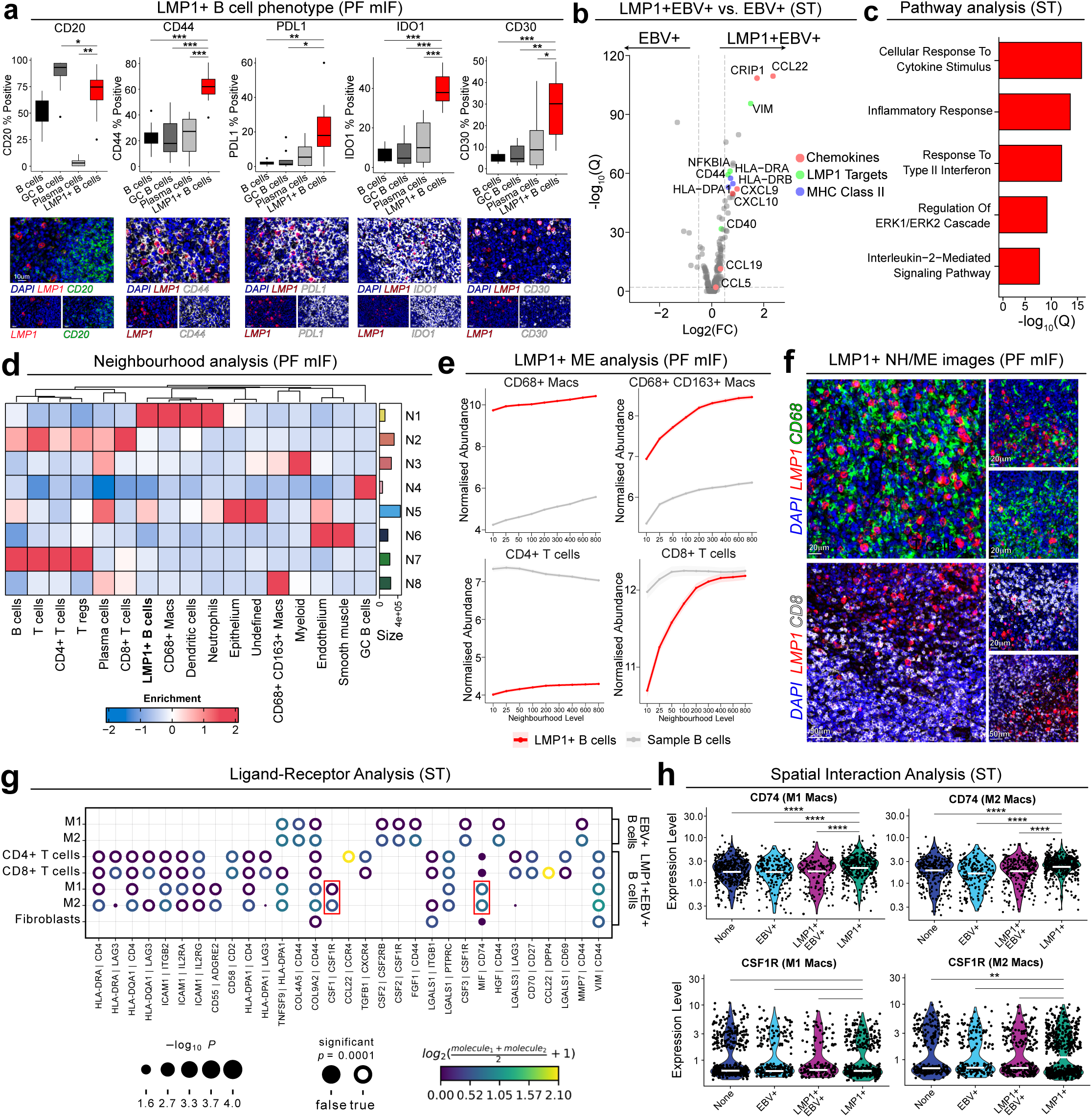
| LMP1 associated chemokines remodel the tonsillar tissue architecture of IM. **a.** Quantification of the phenotype of LMP1+ B cells (PF mIF) showing down-regulation of CD20 (p-value = 0.0499 vs. GC B cells; p-value = 0.0002 vs. plasma cells) and up-regulation of known LMP1 targets CD44 (p-value = 0.0003 vs. B cells; p-value = 0.0006 vs. GC B cells; p-value = 0.0006 vs. plasma cells), PDL1 (p-value = 0.007 vs. B cells; p-value = 0.0207 vs. GC B cells), IDO1 (p-value = 0.0002 vs. B cells; p-value = 0.0002 vs. GC B cells; p-value = 0.0003 vs. plasma cells) and CD30 (p-value = 0.0003 vs. B cells; p-value = 0.0011 vs. GC B cells; p-value = 0.0499 vs. plasma cells). Representative images of co-expression are also included. All comparisons were tested, and non-significant comparisons are not shown. **b.** Volcano plot showing differential expression analysis of LMP1+EBV+ B cells vs. EBV+ B cells from ST data. Some chemokines, LMP1 targets and MHC class II genes are highlighted. **c.** Pathway analysis from ST data of genes upregulated in LMP1+EBV+ B cells vs. EBV+ B cells. **d.** Neighbourhood analysis (see Methods) of IM tonsil tissues from PF mIF data showing the enrichment of cell types in each calculated cellular neighbourhood. **e.** Microenvironment analysis showing how the frequency of CD68+ macrophages, CD68+CD163+ macrophages, CD4+ T cells and CD8+ T cells surrounding LMP1+ B cells (red) and normal B cells (grey) changes as a function of distance. **f.** Representative images (PF mIF) of the enrichment of macrophages (CD68+; green) and depletion of T cells (CD8+; white) around LMP1+ B cells (red). **g.** Ligand-receptor analysis (CellPhone DB; see Methods) of ST data showing the interaction of different immune cell subsets with the EBV+ B cell and LMP1+EBV+ B cell clusters. Red boxes surround interactions of particular interest. **h.** Spatial validation of ligand-receptor analysis showing expression of CD74 and CSF1R on CD68+ and CD68+CD163+ macrophages in the microenvironment of EBV+ B cells and LMP1+EBV+ B cells.

We next wanted to determine if the expression of these immune modulatory molecules was associated with specific microenvironmental features. A neighbourhood analysis of the PF mIF data identified eight distinct tissue neighbourhoods (Fig. 3d & Extended Data Fig. 3a). LMP1+ B cells were enriched in neighbourhood 1, consistently co-located with CD68+ macrophages, which may be due to the increased cytokine and chemokine expression. However, CD4+ and CD8+ T cells were localised to neighbourhoods lacking LMP1+ B cells (Fig. 3d). To assess immune cell infiltration in greater detail, we analysed the immediate microenvironment of LMP1+ B cells by calculating the abundance of cell types in concentric circles of increasing size (see Methods). Compared to EBV-negative B cells, LMP1+ B cells consistently had more macrophages (CD68+ and CD68+CD163+) and fewer CD4+ and CD8+ T cells in their immediate microenvironment (Fig. 3e-f & Extended Data Fig. 3b). We next performed a ligand receptor analysis on the CosMx ST data to identify cognate ligands/receptors pairs that might mediate the recruitment of immune cells, especially macrophages, to the LMP1+EBV+ B cell microenvironment (see Methods). Among the predicted interactions, we found a unique predicted engagement of LMP1+EBV+ B cells with CD68+ and CD68+CD163+ macrophages via the MIF-CD74 and CSF1-CSF1R ligand-receptor pairs (Fig. 3g). MIF (macrophage migration inhibitory factor) binds to CD74 and is a key regulator of macrophage activation, while CSF1 (colony-stimulating factor 1) interacts with CSF1R to promote macrophage survival and differentiation (Fig. 3g). To validate these interactions, we defined niches around EBV+ B cells and LMP1+EBV+ B cells within the CosMx ST data, categorising each cell as belonging to either the EBV+ B cell niche, the LMP1+EBV+ B cell niche, both EBV+ and LMP1+EBV+ B cell niche (EBV+/LMP1+EBV+) or none of the previous, and assessed the expression of chemokine receptors within these niches (Extended Data Fig. 3c). Consistent with the ligand-receptor analysis, we found that within the LMP1+EBV+ B cell niche, CD74 was consistently expressed on CD68+ and CD68+CD163+ macrophages and CSF1R notably upregulated on CD68+CD163+ macrophages (Fig. 3h & Extended Data Fig. 3d). While CXCL9 and CXCL10 were upregulated on LMP1+EBV+ B cells, the cognate ligand CXCR3 was significantly downregulated on infiltrating CD68+ and CD68+CD163+ macrophages (Extended Data Fig. 3e). However, internalisation and downregulation of CXCR3 after ligand binding at the RNA and protein level may explain this^34^. This phenomenon was also present on the low numbers of CD4+ T cells that infiltrated LMP1+ B cell rich zones (Extended Data Fig. 3e). Taken together, these data indicate that CXCL9, CXCL10 and MIF expression on LMP1 expressing EBV-infected cells drives the recruitment of both pro- and anti-inflammatory macrophages.

### PDL1+IDO1+ macrophages form a restrictive niche around LMP1+ B cells

We further interrogated the PF mIF data to analyse the phenotype of macrophages in the LMP1+ B cell niche in more detail. CD68+ macrophages in this niche showed enrichment of PDL1 and IDO1 expression (Fig. 4a & Extended Data Fig. 4a). PDL1+IDO1+ CD68+ macrophages were also significantly more frequent in the microenvironment of LMP1+ B cells compared to EBV-negative B cells (Fig. 4b & Extended Data Fig. 4c). While PDL1+IDO1+ CD68+CD163+ macrophages were most abundant in neighbourhood 8, they were also enriched in the immediate vicinity of LMP1+ B cells (Extended Data Fig. 4b). Using the CosMx ST data, we re-clustered macrophage population, identifying seven distinct functional states (Fig. 4d) defined as CCL18+, SPP1+, IL1B+, CXCL10+, MT+, IDO1+, collagen-producing, and one state without a clear functional signature. Spatial analysis revealed that IDO1+, CXCL10+, and SPP1+ macrophages were specifically enriched in the immediate LMP1+EBV+ B cell niche (Fig. 4e). In contrast, IL1B+ and CCL18+ macrophages were in distinct regions, separate from LMP1+EBV+ or EBV+ B cells. While PDL1 (encoded by CD274) did not define a specific macrophage cluster (Extended Data Fig. 4c), it was significantly overexpressed in LMP1+EBV+ B cell niches compared to other niches (Supplementary Table 7). Differential gene expression analysis of these macrophage subsets revealed that the SPP1+ population exhibited a suppressive phenotype, characterised by the expression of CHI3L1, MMP9, GPNMB (Extended Data Fig. 4d & Supplementary Table 8)^35,36^. Notably, this population also overexpressed MMP12, which cleaves and inhibits IFNG^37^. In contrast, IDO1+ macrophages were enriched for genes involved in antigen presentation (including multiple HLA genes, TAP1, TAP2, and CIITA), while CXCL10+ macrophages showed increased expression of chemokine-related genes linked to TLR signalling (Extended Data Fig. 4e-f & Supplementary Tables 9 & 10). Differential gene expression analysis comparing macrophages in the LMP1+EBV+ B cell niche to those in other niches revealed enrichment of antigen presentation genes (Fig. 4f & Supplementary Table 7), a finding that was also reflected in the PF mIF data (Fig. 4g).

**Figure 4.**
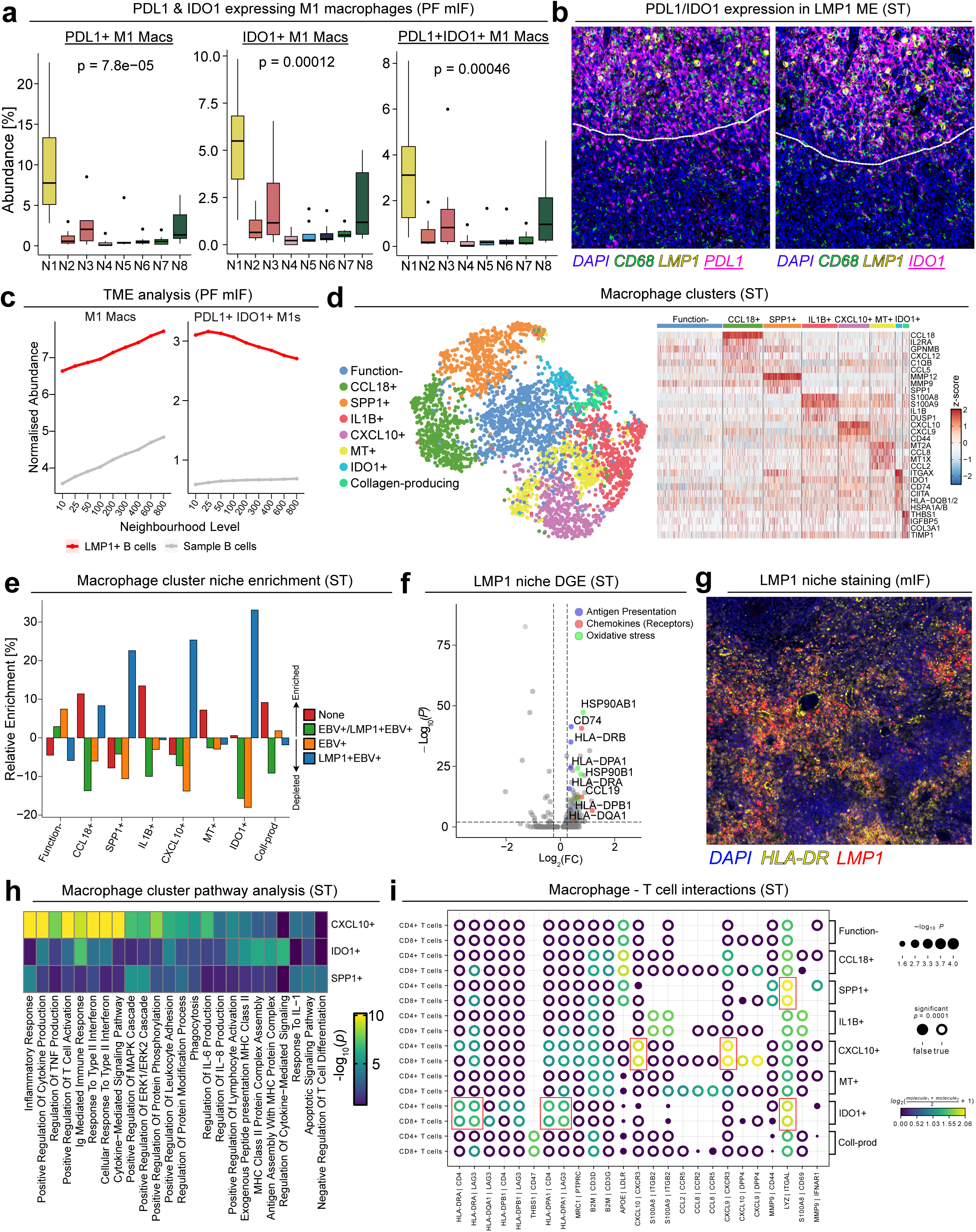
| PDL1+IDO1+ macrophages form a restrictive niche around LMP1+ B cells. **a.** Percentage of PDL1+, IDO1+ and PDL1+IDO1+ CD68+ macrophages as a percentage of all cells per cellular neighbourhood (statistical test: Kruskal-Wallis). **b.** Representative images (PF mIF) of PDL1 (purple; left image) and IDO1 (purple; right image) co-expressing with the macrophage marker CD68 (green) localised to LMP1 (yellow) rich regions. **c.** Analysis of the CD68+ macrophage and PDL1+IDO1+ CD68+ macrophage phenotype frequency as a function of distance from LMP1+ B cells (red) vs. normal B cells (grey). **d.** Re-clustering of macrophages from ST data identified eight sub-clusters defined by their expression profile shown in the heatmap of top differentially expressed genes. **e.** Relative enrichment/depletion of each macrophage sub-cluster present in the niche of only LMP1+EBV+ B cells, only EBV+ B cells, both LMP1+EBV+ and EBV+ B cells (EBV+ /LMP1+EBV+) and neither (None). **f.** Differential gene expression of macrophages located in the LMP1+EBV+ B cell niche vs. other macrophages highlighting antigen presentation (blue), chemokine (red) and oxidative stress (green) genes. **g.** Representative images (PF mIF) of MHC Class II expression on macrophages located in the immediate niche of LMP1+ B cells. **h.** Pathway analysis of genes overexpressed in LMP1+EBV+ B cell niche enriched macrophage sub-clusters (CXCL10+, IDO1+ and SPP1+). **i.** Ligand-receptor analysis (CellPhone DB; see Methods) of ST data showing the interaction of macrophage sub-clusters with CD8+ and CD4+ T cells. Red boxes surround interactions of particular interest.

Pathway analysis indicated that the CXCL10+ macrophage cluster were likely to be pro-inflammatory and involved in cytokine production and T cell activation (Fig. 4h). This macrophage cluster also exhibited evidence of cellular stress, indicated by the upregulation of heat shock proteins (HSP90B1, HSPB1, and HSP90AB1). Notably, large necrotic regions were observed in 3 out of 8 tissue cores, and these regions were enriched for LMP1+ B cells (Extended Data Figure 4g); it is possible the inflammatory environment created by CXCL10+ macrophages are responsible, at least in part, for this tissue damage. In the pathway analysis we also observed that the IDO1+ macrophage cluster were enriched for genes associated with MHC Class II presentation and the SPP1+ macrophage cluster with genes involved in the negative regulation of T cell differentiation (Fig. 4h). This observation prompted us to explore the possibility that these two subsets might interact with T cells to restrict their infiltration and activity. A ligand-receptor analysis identified IDO1+ macrophages as having the strongest MHC Class II interactions with both CD4+ and CD8+ T cells via LAG3, a key inhibitory receptor whose major ligand is MHC Class II (Fig. 4I). Additionally, CXCL10+ macrophages were predicted to engage CD4+ and CD8+ T cells through the CXCL9/10-CXCR3 axis, which regulates T cell recruitment. Taken together these findings suggest that macrophages play a crucial role in controlling T cell infiltration and function in the LMP1+EBV+ B cell microenvironment.

### CD8+ T cells in LMP1+ B cell niches exhibit an activated phenotype

Next, we examined the phenotype and spatial distribution of T cell populations in greater detail. While previous studies of tissue from IM patients have reported a balanced CD8+:CD4+ T cell ratio^21^, our RNA and protein analysis revealed an overall enrichment of CD8+ T cells (Extended Data Fig. 5a-b). However, despite this increase, CD8+ T cells are apparently not able to effectively control EBV-infected cells, suggesting their function may be compromised. To explore this, we re-clustered the PF mIF data focusing on the expression of inhibitory immune checkpoints. Almost one-half of the CD8+ T cells expressed at least one of PD1, LAG3, or TIM3 (Fig. 5a). PD1+ CD8+ T cells were more frequent in the microenvironment of EBV-negative cells, whereas LAG3+ CD8+ T cells were enriched in the microenvironment of LMP1+ B cells (Fig. 5b). No T cell subsets exhibited high Ki67 expression and only the LAG3+TIM3+ and LAG3+TIM3+PD1+ clusters (11.5% of total CD8+ T cells) expressed granzyme B. Thus, in IM tissues, CD8+ T cells expressing inhibitory markers appear to be characterised by diminished cytotoxic and proliferative activity (Extended Data Fig. 5c).

**Figure 5.**
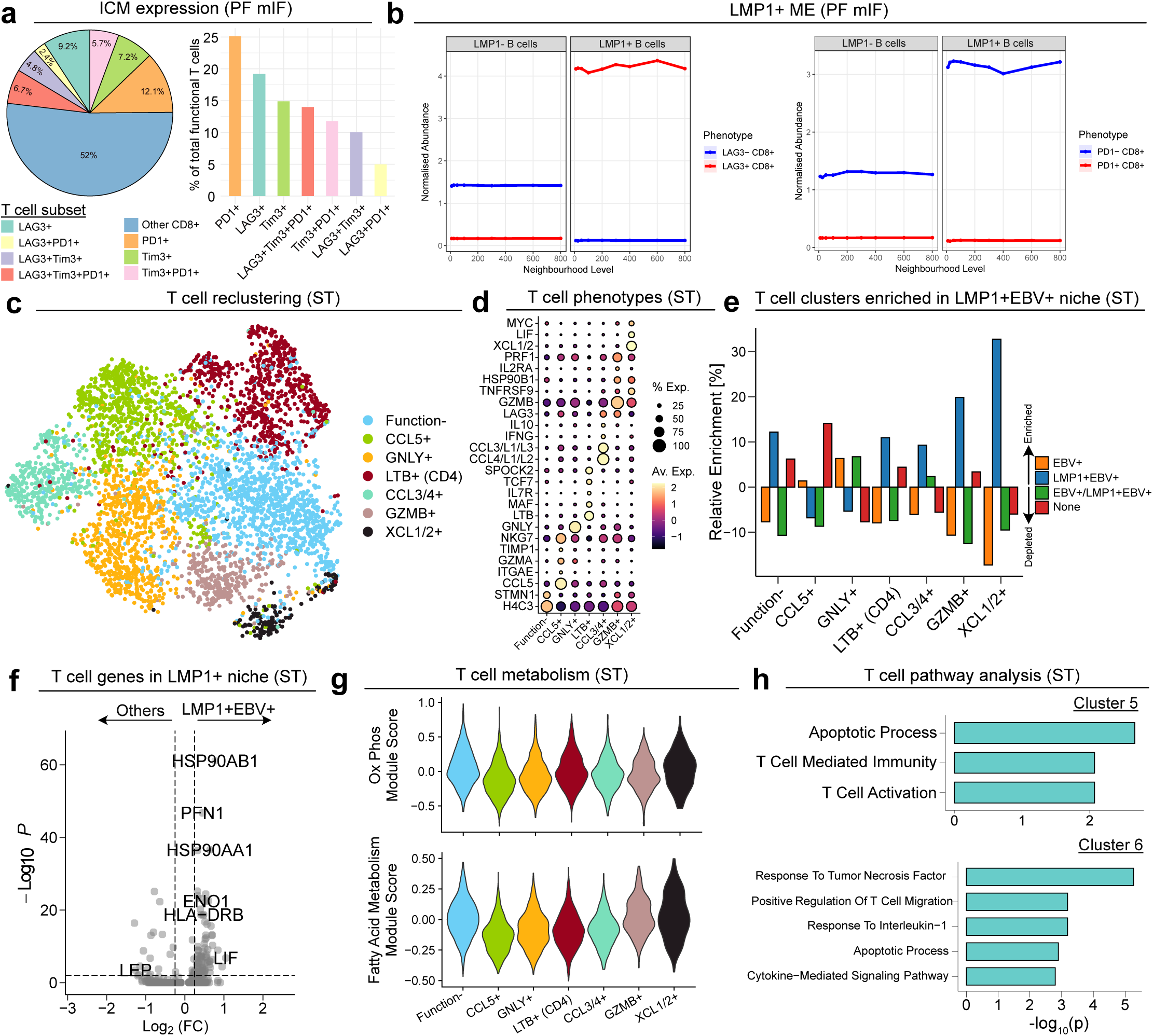
| CD8+ T cells in LMP1+ B cell niches exhibit a chronically active phenotype. **a.** Relative abundance of the expression of immune checkpoint molecules (ICM) PD1, LAG3 and TIM3 on CD8+ T cells by PF mIF. **b.** Abundance of LAG3+ and PD1+ CD8+ T cells in the microenvironment of LMP1+ B cells by PF mIF. **c-d.** Re-clustering of T cells from ST data identified seven sub-clusters defined by their expression profile shown in **D.** the dot plot of top differentially expressed genes. **e.** Relative enrichment/depletion of each T cell sub-cluster present in the niche of only LMP1+EBV+ B cells, only EBV+ B cells, both LMP1+ and EBV+ B cells (EBV+/LMP1+EBV+) and neither (None). **f.** Differential gene expression of T cells located in the LMP1+ B cell niche vs. other T cells. **g.** Expression of metabolism related gene sets in T cell subsets of ST data. **h.** Gene ontology pathway analysis of genes overexpressed in GZMB+ and XCL1/2+ T cells (T cells enriched in LMP1+ B cell niches).

To gain further insights into the function of T cell populations infiltrating the LMP1+ B cell niche, we re-clustered the T cells from the ST dataset into six subsets of CD8+ T cells (Fig. 5c-d & Extended Data Fig. 5d) characterised by high expression of the cytotoxic effector molecules GNLY or GZMB; or the chemokines CCL5, CCL3/4 or XCL1/2; and a subset lacking functional identity. A single subset of CD4+ T cells expressed high levels of LTB (Fig. 5d). We calculated the extent to which each of these T cell clusters was either enriched or depleted within the different cellular niches. Although the GZMB+ and XCL1/2+ clusters contained the fewest numbers of cells, they were most enriched in the LMP1+EBV+ B cell niche (Fig. 5e). CD8+ T cells expressing GNLY, an antimicrobial molecule present in cytotoxic granules of CD8+ T cells, were the only T cell cluster enriched within the EBV+ B cell niche^38^. Differential expression analysis of GZMB+ CD8+ T cells compared to other CD8+ subsets revealed upregulation of GZMH, PRF1, and CD137 (TNFRSF9), and high expression of heat shock proteins (HSP90AB1, HSP90AA1, and HSPB1) (Extended Data Fig. 5e & Supplementary Table 11)^39^. These HSPs as well as CD137 were also expressed in the XCL1/2+ CD8+ T cell cluster (Extended Data Fig. 5f & Supplementary Table 12). T cells within the LMP1+ niche also showed increased expression of these HSPs along with the inhibitory marker PFN1^40^ (Fig. 5f & Supplementary Table 13).

Given the abundance of activated macrophages in the LMP1+EBV+ B cell niche, we investigated whether T cell HSP expression might be driven by increased metabolic demands. Calculating the expression of metabolism gene sets that were present in the ST panel, we found that XCL1/2+ T cells showed the highest expression of genes involved in oxidative phosphorylation and fatty acid metabolism, but not glycolysis (Extended Data Fig. 5g). This cluster also showed the highest expression of inflammatory response genes (Extended Data Fig. 5g). We then identified enriched pathways in the overexpressed genes of the XCL1/2+ and GZMB+ T cell clusters. XCL1/2+ T cells exhibited a strong TNF response, which may promote T cell activation/proliferation or apoptosis^41^. Similarly, while GZMB+ T cells expressed cytotoxic molecules, they were also enriched for genes associated with apoptosis (Fig. 5h). T cells within the LMP1+ B cell niche expressed TIGIT, HAVCR2 (TIM3), ICOS, and TCF7 (Extended Data Fig. 5h) but showed no RNA-level expression of PDCD1 (PD1), TOX or LAG3, indicating a distinct checkpoint profile associated with T cell activation. Together, our data suggest that CD8+ T cells infiltrating the LMP1+ B cell niche have dampened effector functions, express inhibitory immune checkpoints and exhibit gene expression profiles consistent with cellular apoptosis.

## Discussion

In this study, we present the first comprehensive multiomic spatial analysis of tonsillar tissues from patients suffering from infectious mononucleosis, and in doing so provide novel insights into the *in vivo* dynamics of primary EBV infection. Our findings challenge existing models of EBV latency and immune interactions, detailing for the first time with high-plex analysis, the interplay between EBV-infected B cells, epithelial cells, and the immune microenvironment. A key observation from our study is the rare but definitive presence of EBV-infected epithelial cells in IM tonsil tissues. While previous studies have identified EBV-positive epithelial cells in asymptomatic carriers and immunosuppressed individuals^17,42,43^, our data provide the first robust evidence of EBV infection in epithelial cells during primary symptomatic infection. Using spatial transcriptomics, we found that these EBV-positive epithelial cells predominantly expressed markers of superficial squamous differentiation, such as CD44, and displayed upregulated immunoglobulin genes (IgG1/2) in the absence of B cell lineage markers. This observation aligns with recent reports of an epithelial-to-mesenchymal transition (EMT) phenotype in EBV-driven epithelial malignancies, suggesting that EBV may modulate epithelial plasticity during primary infection. The localisation of EBV-positive epithelial cells within tonsillar crypts supports a model in which epithelial cells act as intermediaries in EBV transmission, facilitating viral entry and exit between the oropharynx and the lymphoid compartment.

Our integrated multiomics approach also allowed us to comprehensively phenotype viral and cellular gene expression in the infected B cell populations and to explore the extent to which our findings aligned with existing models of B cell infection. The most widely accepted, so-called, germinal centre model is based mainly on data from highly sensitive PCR assays of whole blood and single-plex immunohistochemistry. In this model, newly infected B cells express a latency III program which drives their proliferation. Subsequently, EBV-infected cells enter a germinal centre reaction and express a latency II program characterised by co-expression of the latent membrane proteins, LMP1 and LMP2. Infected cells then emerge from the germinal centre to differentiate into plasma cells or memory B cells, the latter largely silencing EBV gene expression. While latency III has long been considered a dominant program in newly infected B cells, our findings indicate that this phenotype is exceedingly rare in IM tonsil tissues. Instead, we observed a mutually exclusive pattern of EBNA2 and LMP1 expression, with EBNA2-expressing cells predominantly localised near the tonsillar epithelium and LMP1-expressing cells residing deeper in the lymphoid stroma, in keeping with previous low-plex immunohistochemistry findings^44^. This suggests that primary infection follows a temporal sequence in which EBNA2-positive cells, potentially representing newly infected B cells, transition to an LMP1-positive state as they migrate further into the tissue. However, in keeping with previous studies LMP1 and LMP2A were rarely co-expressed, and LMP1-positive cells were absent from germinal centres^44^. We also observed mutually exclusive expression of BZLF1 and LMP2A, which is in line with previous studies showing that LMP2A suppresses the EBV lytic cycle in B cells^28,29^. Taken together, our data challenge the traditional model of EBV latency in primary infection.

Our data further reveal that LMP1-positive B cells orchestrate a profound remodelling of the local immune microenvironment through the secretion of chemokines such as CXCL9, CXCL10, MIF and CSF1. These chemokines drive the recruitment and activation of macrophages, previously shown at the tissue level in primary EBV infection^45^, which in turn may restrict T cell infiltration. EBV BARF1 is a CSF1 antagonist^46^ that we show is differentially overexpressed in EBV+ compared to LMP1+ B cell clusters, potentially explaining the difference in macrophage abundances in their microenvironments. Spatial analysis showed that LMP1+ B cells were surrounded by a niche enriched for CD68+ macrophages expressing IDO1 and PDL1, known immunosuppressive molecules that contribute to T cell dysfunction^47–49^. This macrophage-rich environment was associated with the depletion of both CD4+ and CD8+ T cells, reinforcing the notion that EBV-infected B cells actively shape an immune-privileged niche that facilitates viral persistence.

Despite the restricted infiltration of T cells into LMP1+ niches, our findings indicate that those CD8+ T cells that are present in this niche retain cytotoxic potential but express many immune checkpoint markers. Almost one-half of the CD8+ T cells expressed at least one inhibitory immune checkpoint molecule, in keeping with data from the blood of IM patients^50^, such as PD1, LAG3, or TIM3, with LAG3+TIM3+ T cells preferentially localised near LMP1+ B cells. These cells also displayed a transcriptional signature associated with chronic activation and metabolic stress. The presence of TIGIT+, ICOS+ and TCF1+ T cells in LMP1+ niches may further support the hypothesis of chronic antigenic stimulation in primary infection.

Collectively, these findings significantly advance our understanding of EBV pathogenesis by providing an integrated spatial model of *in vivo* primary infection. We propose that EBV initially infects epithelial cells within the tonsillar crypts before spreading to B cells, which enter an EBNA2-dominant state near the epithelium. As these infected B cells migrate deeper into the lymphoid tissue, they transition to express LMP1, in turn acquiring a phenotype that promotes immune evasion through macrophage recruitment with concomitant physical and functional impairment of CD8+ T-cells. This spatial and temporal regulation of EBV infection provides a revised framework for understanding how the virus establishes persistence while evading immune-mediated clearance.

In summary, our spatial characterisation provides the first annotated accessible data-rich resource for the study of primary symptomatic EBV infection of palatine tonsils. By integrating spatial transcriptomics and multiplex immunofluorescence, we reveal a complex interplay between EBV-infected cells and the immune microenvironment that shapes the outcome of infection. These insights not only refine our understanding of EBV latency and immune evasion but also highlight potential targets for therapeutic intervention aimed at disrupting the viral niche and enhancing immune control.

## Extended Data Figure Legends

**Extended Data Figure 1.**
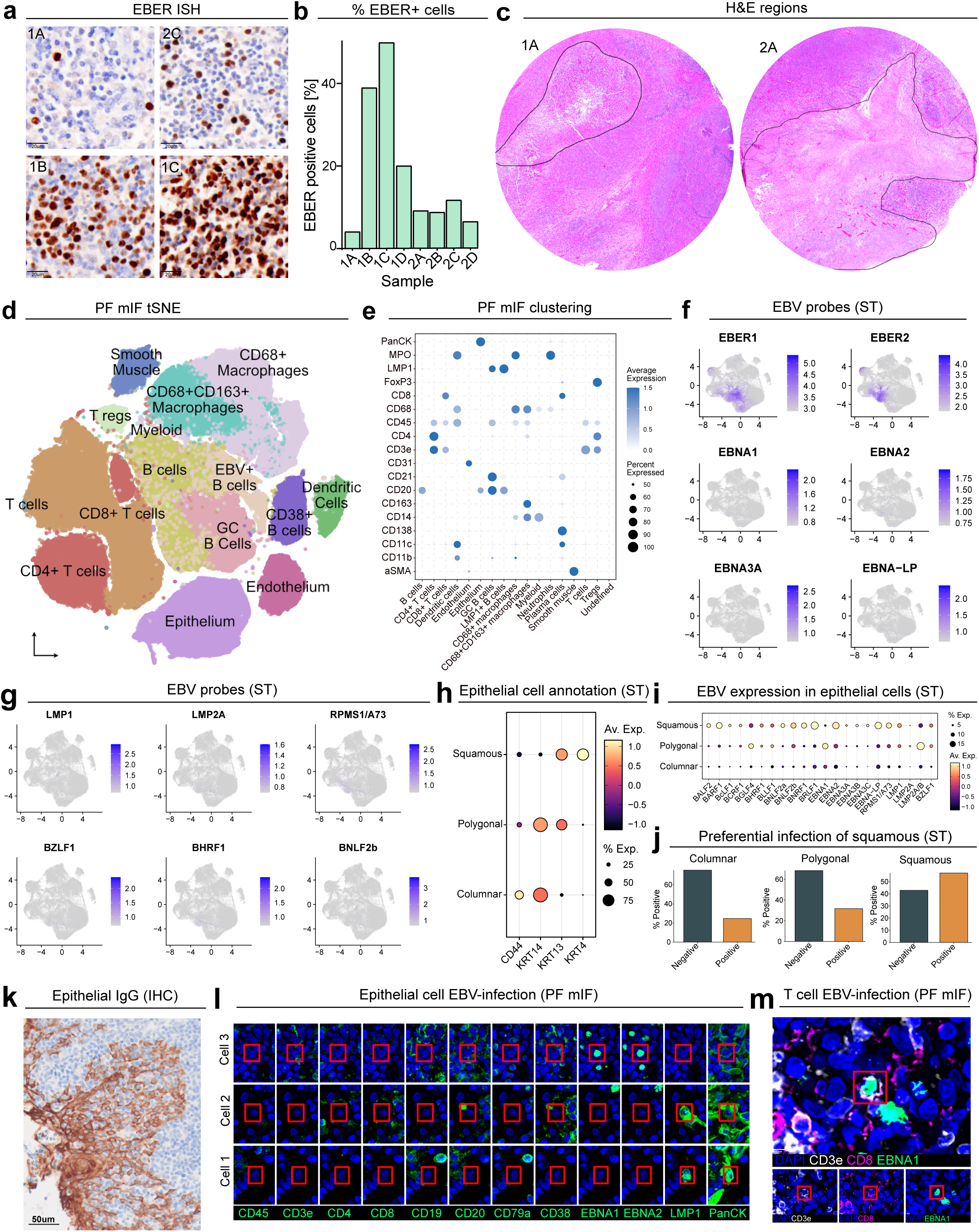
| EBV infects epithelial cells in the tonsil. **a.** Representative images of four IM tonsil samples showing variable EBER expression. **b.** EBER quantification (as percentage positive of total cells) of all IM tonsil samples. **c.** Representative H&E region selection to remove areas of necrosis for downstream analysis. **d.** tSNE plot of PF mIF clustering showing the 15 annotated cell clusters. **e.** PF mIF dot plot showing expression of lineage markers which defined the main annotated cell clusters from D. **f-g.** Feature plot UMAP showing EBV gene expression from ST data in all cell types for **F.** EBER1, EBER2, EBNA1, EBNA2, EBNA3A, EBNA-LP and **G.** LMP1, LMP2A, RPMS1/A73, BZLF1, BHRF1 and BNLF2b. **h.** Dot plot of CD44, KRT14, KRT13 and KRT4 expression from ST data used to annotate the epithelial cell clusters as either squamous (superficial), polygonal (intermediate) or columnar (basal). **i.** EBV gene expression patterns in epithelial cell clusters from ST data. **j.** Preferential infection of squamous (superficial) epithelial layer by EBV (defined by positive expression of EBER1 and EBER2). **k.** Expression of sialylated immunoglobulin (RP215) in the epithelium of IM tonsil samples. **l.** PF mIF image panels showing co-expression of LMP1 and pan-cytokeratin (cell 1), EBNA1, EBNA2 and pan-cytokeratin (cells 2 and 3), and lack of expression of immune cell markers, CD45, CD3e, CD4, CD8, CD19, CD20, CD79a and CD38. All images are pseudo-coloured green. **m.** T cell infection showing co-expression of CD3e, CD8 and EBNA1.

**Extended Data Figure 2.**
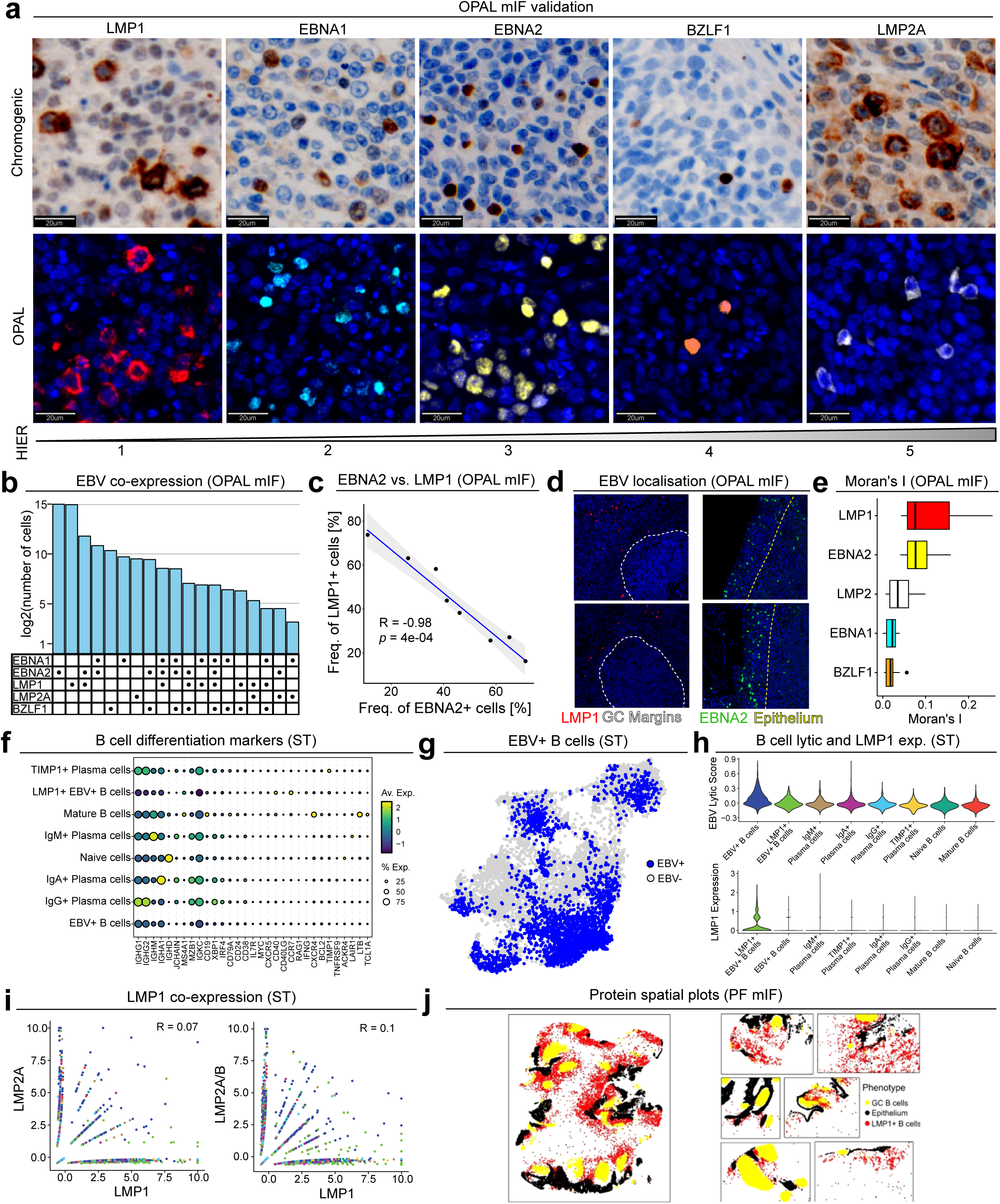
| Spatio-temporal evolution of EBV-infection in tonsillar tissue. **a.** OPAL mIF detection of LMP1, EBNA1, EBNA2, BZLF1, and LMP2A EBV proteins was validated through a two-step process. Initially, antibodies were assessed using chromogenic detection (DAB) at the appropriate antigen retrieval conditions, followed by fluorescent validation with their corresponding fluorophores. **b.** Frequency of cells expressing different EBV co-expression patterns. **c.** Correlation between the frequency of EBNA2+ cells and LMP1+ cells in each IM tonsil (OPAL mIF). **d.** EBV protein localisation of EBNA2+ cells close to epithelial beds and LMP1+ cells to around germinal centres (OPAL mIF; dashed white lines define manually annotated epithelium and germinal centres). **e.** Moran’s I calculation of EBV protein expression cells (from OPAL mIF) showing higher co-localisation of LMP1+ and EBNA2+ cells. **f.** Dot plot showing B and plasma cell differentiation gene expression from ST data. **g.** EBV+ B cells (defined by positive expression of EBER1 and EBER2 ST data) plotted on the B and plasma cell UMAP. **h.** EBV gene expression lytic score and LMP1 expression in each B and plasma cell cluster. **i.** Co-expression of LMP1 and LMP2A/LMP2A/B EBV genes from ST data confirming protein data. **j.** Spatial plots (PF mIF) confirming LMP1+ B cell localisation around germinal centres at the protein level.

**Extended Data Figure 3.**
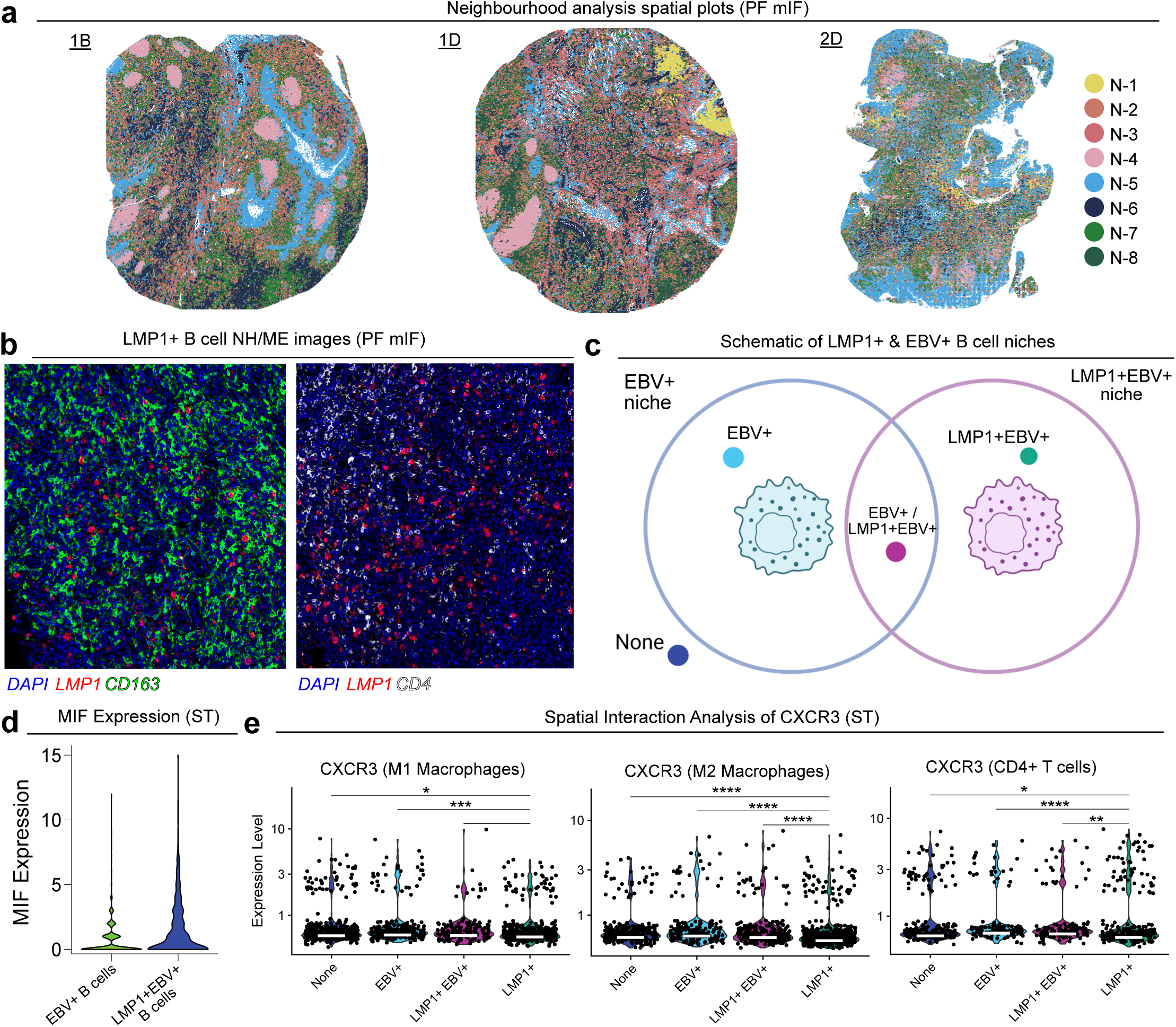
| LMP1 associated changes in IM tissue architecture. **a.** Spatial plots (PF mIF) showing the arrangement of cellular neighbourhoods within three representative IM tonsil tissue cores. **b.** More representative images (PF mIF) of the enrichment of CD68+CD163+ macrophages (CD163+; green) and depletion of T cells (CD4+; white) around LMP1+ B cells (red). **c.** Schematic of niches. Each cell was defined as being in an EBV+ B cell niche, a LMP1+ B cell niche, both (EBV+/LMP1+EBV+) or in neither. **d.** Expression of MIF on EBV+ B cells and LMP1+EBV+ B cells in ST data. **e.** Spatial validation of ligand-receptor analysis showing reduced expression of CXCR3 on CD68+ and CD68+CD163+ macrophages as well as CD4+ T cells in the niche of LMP1+EBV+ B cells.

**Extended Data Figure 4.**
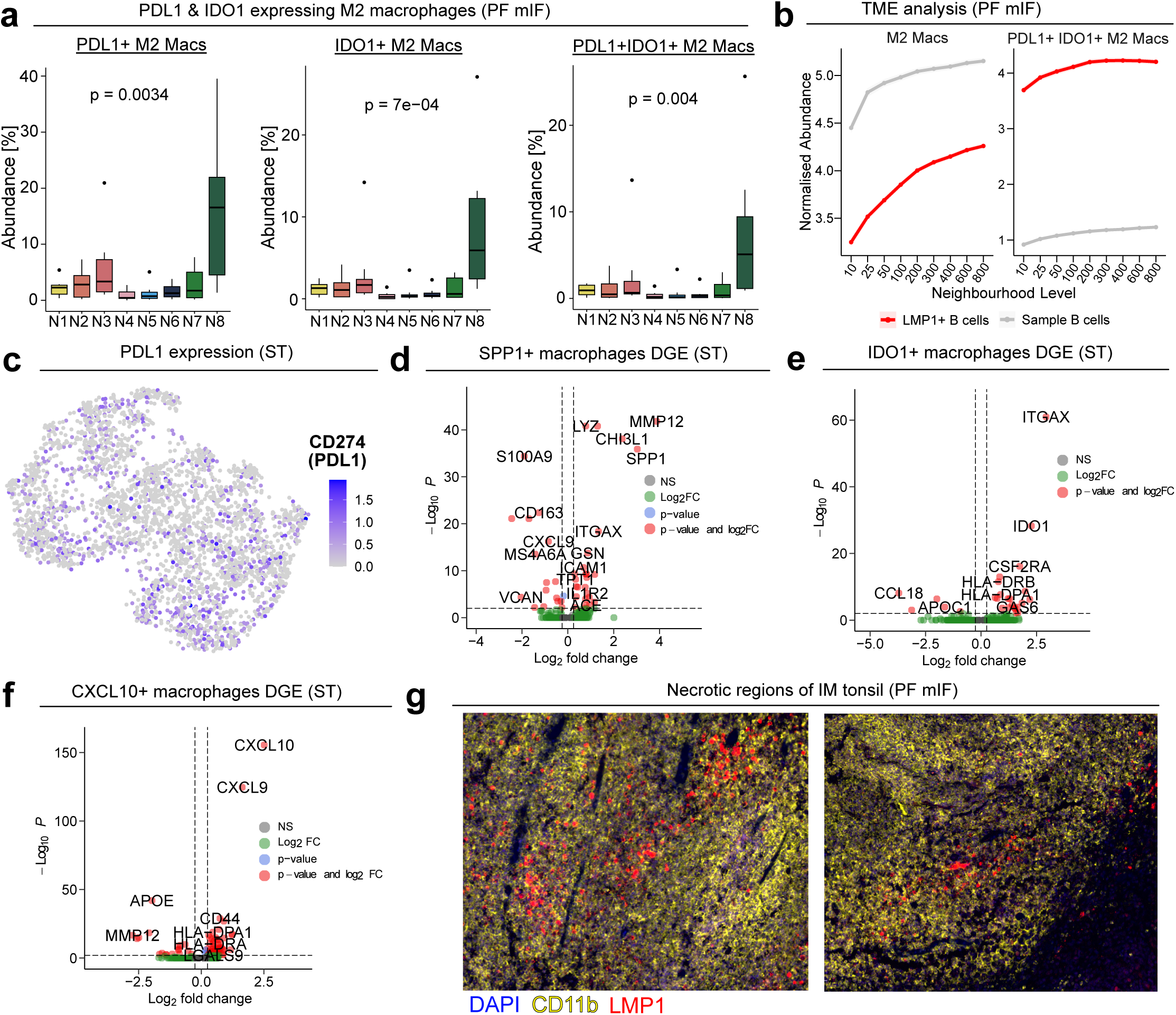
| Macrophage phenotypes in the LMP1+ B cell niche. **a.** Percentage of PDL1+, IDO1+ and PDL1+IDO1+ CD68+CD163+ macrophages as a percentage of all cells per cellular neighbourhood. (statistical test: Kruskal-Wallis). **b.** Analysis of the CD68+CD163+ macrophage and PDL1+IDO1+ CD68+CD163+ macrophage phenotype frequency as a function of distance from LMP1+ B cells (red) vs. normal B cells (grey).**c.** Expression of CD274 (PDL1) on macrophage clusters by ST. **d-f.** Differential expression analysis of **D.** SPP1+, **E.** IDO1+ and **F.** CXLC10+ macrophages vs. all other macrophage clusters. **g.** Representative images (PF mIF) of LMP1+ B cells (red) located within large necrotic regions (denoted by CD11b expression).

**Extended Data Figure 5.**
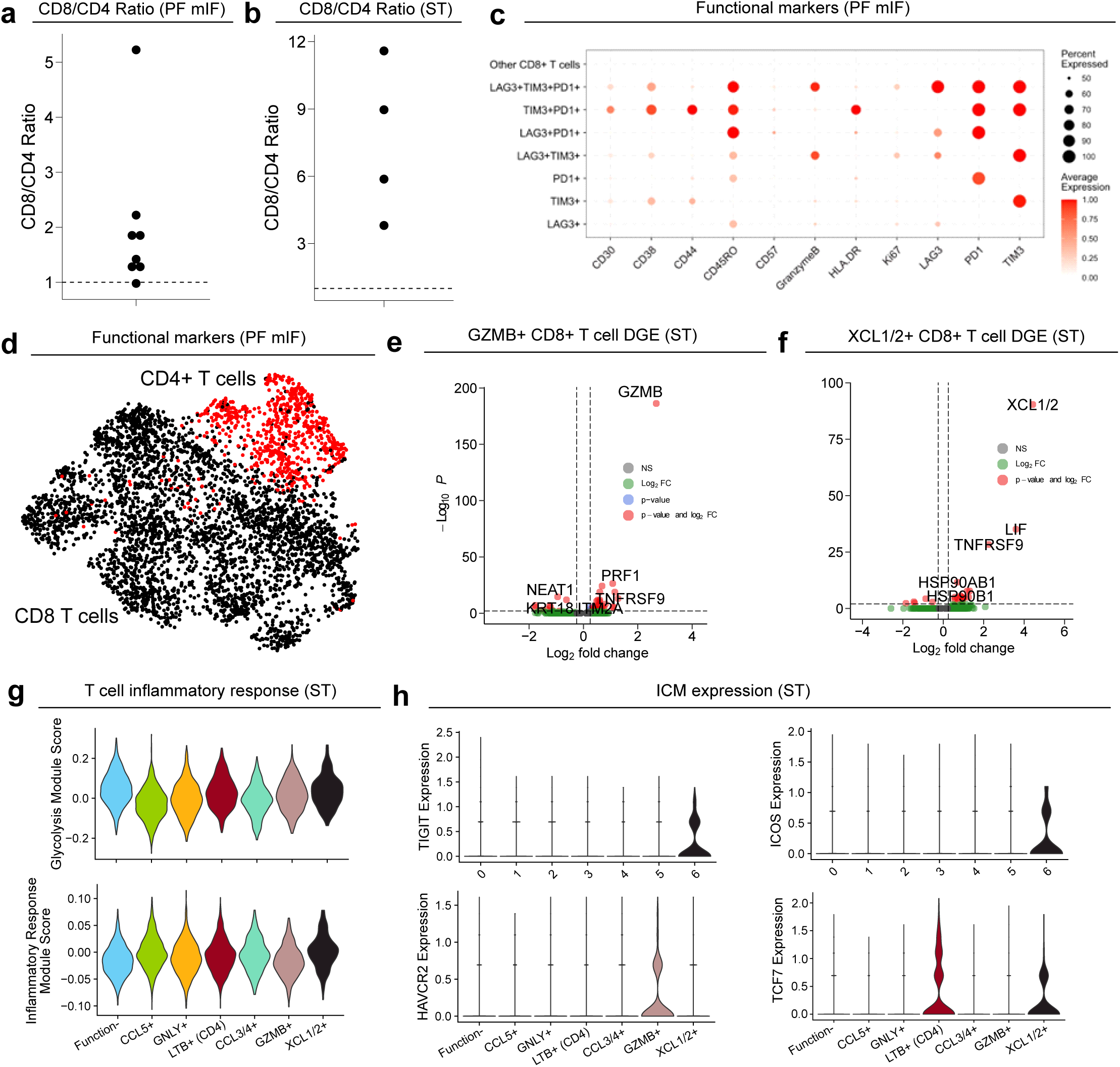
| CD8+ T cell gene expression in LMP1+ B cell niches. **a-b.** CD8 to CD4 T cell ratio of IM tonsil samples in A. PF mIF data and B. the ST data. **c.** Functional marker expression of re-clustered T cells to identify ICM expressing cells. **d.** CD8+ and CD4+ T cells shown on the re-clustered T cell UMAP. **e-f.** Differential gene expression of E. GZMB+ CD8+ T cells and F. XCL1/2+ CD8+ T cells vs. other T cell subsets. **g.** Expression of the inflammatory response gene set within the T cell subsets. **h.** Expression of ICMs LAG3, ICOS, HAVCR2 (encodes TIM3), TIGIT, and PDCD1 (encodes PD1) across the T cell subsets.

## Methods

### TISSUE COLLECTION

#### Tissue cohort & preparation

Seven formalin-fixed paraffin embedded (FFPE) tonsil samples from patients with an infectious mononucleosis (IM) diagnosis (from Prof. Gerald Niedobitek, Sana Klinikum Lichtenberg, Berlin) and a further two FFPE tonsil samples from patients with an IM diagnosis (from Dr Matthew Pugh, University of Birmingham) were obtained as full-face blocks. Tonsillectomies were performed due to severe obstructive tonsillitis. Tissue microarrays (TMA) were constructed from eight IM tonsil samples, two TMAs with four cores/samples each. A 5mm hollow needle was used to punch cores from each IM tonsil blocks in the lesional areas identified by H&E staining. Liver tissue and normal lymph node were placed at the edges of the TMAs for orientation purposes. TMA sections were cut onto positively charged slides for PF mIF, OPAL mIF, CosMx and H&E staining.

### EXPERIMENTAL ANALYSES

#### Haematoxylin and eosin

Tissues were baked for 3 hours prior to staining to ensure tissue adherence. The Leica BondMax H&E protocol F was used to stain haematoxylin and eosin (H&E) before subsequently being mounted with DPX mountant media and scanned (Akoya FUSION).

#### Chromogenic EBER ISH

5μm sections were cut from IM tonsil FFPE blocks. Slides were dewaxed in Histoclear II and rehydrated before target unmasking by boiling in citrate buffer (4.3mM sodium citrate, 1.3mM citric acid, pH 6) for 20mins. Slides were then washed in RNase free water, dipped in 96% EtOH and briefly air dried. A FITC-conjugated EBER probe (Dako) was added and incubated at 55°C for 90mins. Slides were then washed in stringent wash solution (Dako, 15min 55°C) and TBS-T (Tris buffered saline + 0.1% Tween 20). Anti-FITC-alkaline phosphatase conjugated secondary (Dako) was then applied for 35min at room temperature. Following washing with TBS-T, the signal was developed with BCIP (Dako), and counterstained with Vector Red (Vector Labs), dehydrated and mounted with DPX.

#### OPAL mIF optimisation and staining

Each antibody included in the OPAL panel was first optimised manually using chromogenic IHC to ensure their specificity in tissue sections from patients with IM. Antibody concentrations and heat induced epitope retrieval (HIER) buffer were optimised and the stability of the epitopes over subsequent rounds of HIER retrieval was assessed. Epitopes detectable at all HIER rounds were placed later in the panel design whereas epitopes which could not be detected after multiple HIER steps were placed early in the panel to facilitate their detection. In these optimisation experiments, slides were left to cool for 30 minutes before being subjected to the next HIER. Antibodies were tested after 1, 3 and 5 HIERs only. The final antibody clone information, concentrations, panel position, timings and HIER are listed in Table S1.

In brief, the full OPAL panel stainings were completed on the Leica Bond RXm with the following steps. Slides were baked offline for 1 hour in a 60°C oven before loading on the auto-stainer and undergoing the pre-treatment conditions of default bake (30mins, 60°C) and dewaxing. Bond ER 2 buffer (pH 9) was used for HIER at each retrieval step at 95°C. The first HIER step was 40 minutes with all other HIER incubations being 20 minutes. A five-minute endogenous peroxidases blocking step at room temperature with Bloxall was added prior to Opal Blocker for 10 minutes. Primary antibodies were incubated for 1 hour at room temperature. Secondary antibodies were incubated for 30 minutes at room temperature. The Leica secondary antibody was replaced with a Vector Labs secondary for mouse antibodies (EBNA2, LMP1, BZLF1) to increase signal intensity. Abcam anti-rat antibody was used for the detection of rat antibodies (EBNA1, LMP2A). Opal dyes were incubated for either 30 or 45 minutes as depicted in Table S1. Incubations were carried out at room temperature. DAPI was dispensed twice for 5 minutes each. Slides were then imaged on the Akoya FUSION microscope.

#### PhenoCycler-FUSION mIF staining and imaging

A 52-plex PhenoCycler FUSION panel was first designed and validated chromogenically. The panels, clones and barcodes used are available in Table S2. Glass slides were prepared, stained, and fixed as per the PhenoCycler protocol. Briefly, slides were deparaffinised in Xylene and re-hydrated in decreasing concentrations of ethanol (100%, 90%, 70% & 50%). Heat Induced Epitope Retrieval (HIER) was performed in a pressure cooker at 110°C for 18 minutes in Tris-EDTA buffer (pH = 9.0). The tissue was then incubated in PhenoCycler Staining Buffer (Akoya Biosciences) for 30 minutes to block non-specific binding of antibodies. Subsequently, the tissue was incubated in a cocktail of the conjugated antibodies overnight at 4°C, then fixed in 4% paraformaldehyde for 10 minutes, 100% methanol for 5 minutes, Fixative Reagent (Akoya Biosciences) for 20 minutes and stored until imaging. Akoya Reporters were added to the corresponding well of a 96-well plate in preparation for imaging, based on the cycle design of the experiment. Autofluorescence subtraction, stitching and compression were completed in the FUSION software resulting in qptiff files in preparation for processing and analysis.

#### CosMx spatial transcriptomics staining and imaging

The CosMx human universal cell characterisation panel (1000-genes with an additional 24 custom designed EBV-specific probes; Table S5) was applied to 4 FFPE tonsil sample tissues with a mean of 4 fields of view (FOV) per tissue for a total of 16 FOV. B2M, Pan-Cytokeratin, CD45 and CD3 antibodies as well as DAPI were used for visualising cellular morphology. Tissues were prepared, stained and acquired as previously described^20^. Briefly, 5μm TMA sections were cut onto positively charged slides at baked at 60C overnight. Preparation for in-situ hybridization (ISH) was conducted on the Leica Bond RXm by deparaffinization and HIER for 15 minutes at 100 °C using ER2 epitope retrieval buffer (Leica Biosystems, Tris/EDTA, pH 9.0). After HIER, tissue sections were digested with Proteinase K diluted (3µg/ml) in ACD Protease Plus at 40C for 30 minutes, washed twice with diethyl pyrocarbonate (DEPC)-treated water and incubated in 0.00075% fiducials (Bangs Laboratory) in 2x saline sodium citrate, 0.001% Tween-20 (SSCT solution) for 5 minutes at room temperature. Slides were rinsed with 1X PBS, and then fixed with 10% neutral buffered formalin (NBF) for 5 minutes at room temperature before being rinsed twice with Tris-glycine buffer (0.1 M glycine, 0.1 M Tris-base in DEPC H2O) and once with PBS for 5 minutes. Slides were then blocked with 100 mM N-succinimidyl (acetylthio) acetate (NHS-acetate, Thermo Fisher Scientific) in NHS-acetate buffer (0.1 M NaP, 0.1% Tween pH 8 in DEPC H2O) for 15 minutes at room temperature. Sections were rinsed with 2x saline sodium citrate (SSC) for 5 minutes and an Adhesive SecureSeal Hybridization Chamber (Grace Bio-Labs) was placed over the tissue. NanoString CosMx ISH probes were prepared by incubation at 95 °C for 2 min and placed on ice, before the probe mix was pipetted into the hybridization chamber and hybridization was performed at 37C overnight. Tissues were washed twice in 50% formamide in 2x SSC at 37C for 25 minutes, washed twice with 2x SSC for 2 minutes at room temperature, and blocked with 100 mM NHS-acetate in the dark for 15 minutes. Slides were then acquired on the CosMX SMI instrument according to the Nanostring protocol and as previously described^20^. After RNA acquisition, tissues were incubated with a 4-fluorophore-conjugated antibody cocktail against B2M (488 nm), Pan-Cytokeratin (532 nm), CD45 (594 nm), and CD3 (647 nm) with a DAPI counterstain in the CosMx SMI instrument for 2 hours. Reporter Wash Buffer was used to wash any unbound antibodies and Imaging Buffer was added to the flow cell before the five channels were captured on the CosMX SMI machine.

### COMPUTATIONAL ANALYSES

#### OPAL mIF – Image pre-processing

OPAL mIF images were imported into Phenochart and inform for spectral unmixing. We used the default synthetic OPAL library for spectral unmixing in inForm as it was constructed from tonsillar tissue^51^. Additionally, region stamps were taken in Phenochart to fine-tune the autofluorescence (AF) subtraction. The AF dropper tool was used to remove/reduce the AF signal by selecting regions containing areas of high AF, such as red blood cells. Phenochart was then used to select each TMA as a ROI for inForm Batch processing. In inForm, the ROIs were processed using the custom settings defined on the stamps before the component image was exported in tiles. The tiles were stitched into a pyramidal OME-TIFF using QuPath. Images were the processed using a custom image analysis platform which includes tissue de-arraying, marker normalisation, cell segmentation using DeepCell [refs], and marker expression for each channel to obtain cell-protein matrices.

#### OPAL mIF – Image analysis

Guided by H&Es, EBER-ISH and PF mIF staining, images were examined with consultant histopathologist Dr Matthew Pugh to annotate areas unsuitable for analysis which were free from necrotic cells or staining/imaging artefacts. Each cell was denoted positive or negative for each EBV protein by first automatically detecting a cutoff point using MetaCyto before manual adjustment^52^. Cells were then initially assigned to a known latency patterns based off of EBV protein co-expression before also being denoted by their co-expression patterns regardless of latency types. Cell abundances were calculated for each latency type and co-expression pattern. The spatial co-localisation of each marker was calculated using Moran’s I in R v4.2.0. Manual pathological annotation of GCs was conducted by consultant histopathologist Dr Matthew Pugh.

#### PhenoCycler-FUSION – Image pre-processing

Raw qptiff images were processed through CellMAPS^19^. Images were normalised using an automated contrast equaliser. CD45, pan-cytokeratin and CD31 were merged to create a pseudo-membrane marker for segmentation. Segmentation was completed on a per-core basis on the stitched DAPI and pseudo-membrane marker channels using DeepCell^53^. FCS files were generated using the segmentation masks with the ‘regionprops’ function within Python, exporting the average intensity of each channel and morphological features of each cell.

#### PhenoCycler-FUSION – Clustering & annotation

Cell-count matrices were imported into the MISSILe package. Clustering was performed using PhenoGraph^54^. For clustering, lineage markers CD31, CD4, CD163, aSMA, CD68, Pan-CK, CD45, CD138, CD11c, CD8, FoxP3, CD11b, CD14, CD3e, CD56, CD20, LMP1, MPO, Keratin8/18 were used to cluster the cells (k = 70). Clusters which contained markers for more than one cell type were subsetted and re-clustered in a similar manner. Clusters were annotated based on the criteria presented in Table S4. The annotated phenotypes were validated by plotting the cells in their original spatial location and comparing the visual patterns to the original images. To determine thresholds for marker positivity, MetaCyto was run on only functional markers that were not used for cell clustering^55^. EBV proteins were also not included in automatic thresholding as manual thresholds were set to ensure robust identification of EBV-positive cells.

#### PhenoCycler-FUSION – Spatial analysis

To identify recurring patterns of cellular neighbourhoods, we used the algorithm previously described with N=10 and k=10^56^. To identify cells in the immediate vicinity of cells of interest, microenvironment analyses were conducted. We calculated the abundance of specified cell types (e.g. immune cells) in the microenvironment of anchor cells (e.g. EBV-infected cells) in concentric circles of increasing size, namely 25-800 cells. Repeating this analysis with different anchor cells allows for the comparison of the microenvironments of cells of interest (e.g. EBV+ vs. EBV-cells). In some occasions, the population of anchor cells being compared to each other were substantially different in size (e.g. there were much more cells not infected with EBV than infected with EBV in some samples of IM tonsil). In these cases, random samples of cells were selected to compare equal numbers of cells. Moreover, some populations of cells were extremely large (>100,000). These populations were too computationally heavy to run and downsampling was also employed in these instances. All analyses with downsampling were repeated three times to ensure the reproducibility of results.

#### CosMx – Pre-processing

In AtoMx, images were segmented using the antibody-based markers to obtain cell boundaries. Transcripts were then assigned to each cell to obtain a feature-cell matrix. Cells with an average negative control count >0.5 and <20 detected features were filtered out.

#### CosMx – Clustering, validation, etc

Feature-cell matrices were imported into Seurat (v5) [refs]. Clustering of all cells was completed by normalising by SCTransform clipped at -10 and 10, identifying the 800 most variable genes, calculating the PCA with 50 principal components. Neighbouring cells were found with the top 50 dimensions and the clustering resolution was set to 0.5. Cells were the visualised on a UMAP with the top 50 dimensions. Clusters were annotated according to their canonical marker expression using known markers genes as well as differential expression analysis between the clusters using FindAllMarkers. Subsequent re-analysis of individual cell types was completed the same other than setting the number of variable genes to 100.

#### CosMx – Spatial analysis

To investigate the cellular composition and gene expression within the niche of EBV/LMP1 positive cells, we used the RANN to identify the 50 closest cells to each anchor cell type. Cells that were present in multiple anchor cell niches were identified and labelled. Differential gene expression of cells within anchor cell niches were compared and plotted with EnhancedVolcano. Gene pathway analysis was conducted with Enrichr.

#### CosMx – Ligand Receptor analysis

To conduct ligand-receptor analysis, the feature-cell matrix was transformed to an AnnData object and imported in scanpy. CellPhoneDB was ran with nperm=200 and threshold=0.04/0.001 depending on the comparisons.

#### Data visualisation & statistics

All data was plotted in R using ggplot2 and ggpubr (apart from ligand-receptor analysis which was created in Python). mIF images were created in QuPath^57^. Illustrations were created with BioRender.com. The Mann-Whitney U test was used for statistical analyses of unpaired data with non-gaussian distributions unless otherwise specified in the figure legends. P-values of < 0.05 were considered statistically significant throughout.

## Data Availability

Images and numerical data generated for this study will be available upon publication.

## Code Availability

CellMAPS source code can be found at https://www.github.com/EannaFennell/CellMAPS.

## Acknowledgements

C.I.L. is funded by an Irish Research Council Postgraduate Scholarship (GOIPG/2020/142). É.F. is funded by an Irish Research Council Postdoctoral Fellowship (GOIPD/2022/97). This work was partly funded by the Limerick Digital Cancer Research Centre (University of Limerick). All illustrations were created with BioRender.com. The authors would like to acknowledge Nadezhda Nikulina, Oliver Braubach and Akoya Biosciences for imaging full-face IM tonsil sections to assist in the validation and optimisation of antibodies. The authors would also like to acknowledge the Birmingham Tissue Analytics (BTA) facility, Propath UK and Bruker Spatial Biology for assistance in generating the Nanostring CosMx spatial transcriptomic data.

## Author Contributions

C.I.L., P.G.M., & É.F. conceived and designed the study;

M.P., S.D. & G.N. obtained ethical approval for the tissue samples;

C.I.L. & M.P. prepared the samples for staining;

A.H. validated and acquired OPAL mIF images;

C.I.L & N.N. acquired the PhenoCycler FUSION images;

G.S.T. & M.P. developed the CosMx custom EBV probe panel.

M.P. & G.S.T. acquired the CosMx spatial transcriptomics data;

C.I.L. & É.F. analysed the data;

C.I.L. & É.F. prepared the figures;

C.I.L., P.G.M., & É.F. wrote the manuscript;

P.G.M. & É.F. supervised the work.

All the authors contributed to the final version of the manuscript.

